# Good dog, bad wolf: Debunking evolutionary preparedness to fear wolves

**DOI:** 10.64898/2026.01.06.697971

**Authors:** Magdalena Boch, Friederike Range, Sarah Marshall-Pescini, Ronald Sladky, Claus Lamm

## Abstract

The return of wolves to historical ranges often causes strong opposition, contrasting with widespread affection for domestic dogs. Evolutionary accounts suggest this discrepancy stems from deeply ingrained affective responses that predispose humans to fear wolves while fostering positive attachment to dogs. Using neuroimaging and self-report data from 42 participants, we investigated how defensive neural circuits linked to implicit threat detection are related to explicit threat evaluations of the two species. During functional MRI, participants viewed images of wolves, dogs, and snakes - the latter chosen for their strong engagement of predisposed threat-related responses - displaying either aggressive or non-aggressive behaviours. As predicted, aggressive displays of all species elicited stronger neural responses in threat-related brain areas. Snakes elicited stronger subcortical threat responses than wolves or dogs, while neural responses to wolves and dogs were largely indistinguishable when comparing aggressive displays. However, explicit threat evaluations strongly differed: wolves were perceived and rated as significantly more threatening than dogs, especially when the species exhibited aggressive displays. Together, these findings challenge accounts that humans are predisposed for negative responses to wolves, suggesting instead that our attitudes towards the species are shaped primarily by deliberate-evaluative processes rather than by implicit-automatic threat responses. This underscores the role of experience, knowledge, stereotypes, and socio-cultural mechanisms in shaping public perceptions of human-wildlife conflicts and conservation efforts. These results provide new insights into the cognitive and affective foundations of human–animal relationships, informing efforts to improve human-wildlife coexistence.

**Significance statement:** Why are wolves feared while dogs are loved? One common explanation is that humans are evolutionarily predisposed to fear wolves as dangerous predators, while dogs, due to domestication, occupy a special niche in human lives. By comparing neural and behavioural responses to wolves, dogs and snakes, we show that wolves do not evoke stronger threat-related brain responses than dogs, despite being judged as the greater threat. However, wolves and dogs alike engage cortical systems associated with perception and evaluation, suggesting that cultural narratives and explicit reasoning, rather than ‘hardwired’ automatic fear, underlie wolf aversion. These findings challenge evolutionary accounts of fear preparedness and suggest that negative attitudes towards wolves can be addressed through education, communication, and public engagement.

## Introduction

Wolves have long occupied a paradoxical place in human culture. Revered in some traditions and reviled in others, they appear through folklore as cunning predators and regularly stand at the centre of public debates regarding their conservation and return to historical ranges in Europe and America. Wolves often evoke adverse emotional reactions, fuelling opposition to their protection, which likely also influenced the recent downgrading of their protection status by the European Union (1). Negative attitudes towards wolves and their link to sociocultural, demographic, and economic factors have been extensively studied across stakeholder groups (see e.g., 2, 3). However, far less is known about what shapes these attitudes, a knowledge gap that prevents a better understanding of why intervention efforts often fail or even backfire (for a recent Perspective, see 4). In striking contrast, domestic dogs - wolves’ closest living relatives - are widely kept as companions, and human-dog relationships are often characterised by strong affiliative bonds (5–7), despite in reality posing a much greater risk of human injuries and fatalities than wolves due to their numbers and cohabitation (8, 9).

One explanation for the stark divergence in attitudes toward wolves and dogs lies in the dual nature of the perceptual and evaluative processes that shape human attitudes and, in turn, influence behaviour towards animals (4, 10). Implicit processes are strongly rooted in associative learning or evolutionary predispositions influencing attitudes through largely automatic, affect-centred reactions operating outside conscious awareness. Explicit processes, in contrast, involve deliberate, conscious evaluations shaped by cultural norms, beliefs, knowledge and personal experience. While explicit attitudes are amenable to change, e.g., in response to new information or deliberation, implicit processes are more resistant and can strongly influence behaviour outside one’s awareness (4, 10). For instance, information campaigns highlighting that the threat posed by wolves to humans is greatly overestimated (8, 11) may influence individual and public opinions; if met by deeply rooted fears, however, people may still vote against wolf protection measures.

It has been proposed that such fears are the result of a broader evolutionary predisposition to avoid animals posing a severe threat to humans (evolutionary preparedness hypothesis; 12), which has been supported by high prevalences and rapid acquisition of fear, for example, toward snakes or spiders (13– 15). This mechanism may also extend to wolves and other large predators. Interestingly, though, it does not appear to shape our attitudes towards dogs despite their similarity to wolves. This may be an effect of their domestication, replacing threat perceptions with widespread affection and bonding. However, the interplay between implicit and explicit processes in shaping our attitudes towards “bad wolves and good dogs” remains poorly understood (4, 16).

While explicit attitudes can be readily assessed through self-report, implicit processes are more difficult to measure, and behavioural measures of implicit bias have recently been questioned (17). Accordingly, we used functional magnetic resonance imaging (fMRI) to examine neural responses to threatening images of wolves and dogs, using snakes as an evolutionarily salient reference species for which preparedness to fear has been repeatedly demonstrated (15, 18, 19). Our approach was guided by recent dual-systems accounts of fear and anxiety (20), distinguishing subcortical circuits implicated in automatic threat detection from neocortical regions associated with conscious evaluation and emotional awareness. To identify threat-related activation, we included non-aggressive (neutral) displays of each species, allowing us to contrast neural responses to threatening cues against less threatening baselines.

We predicted that if negative attitudes toward wolves reflect an implicit, evolutionarily grounded threat bias, wolves would elicit stronger activation in brain regions associated with defensive threat encoding than dogs, and this activation would be more similar to that evoked by snakes. Self-reported threat ratings were used to capture explicit attitudes and evaluative processes, with wolves and snakes predicted to be perceived and evaluated as more threatening than dogs. If the neural responses to wolves, however, were to resemble those to dogs rather than to snakes, this would challenge accounts positing unaware and automatic mechanisms as the primary drivers of negative attitudes toward wolves.

## Results

While undergoing functional MRI, 42 participants were presented with images of dogs, wolves and snakes displaying either a threatening (aggressive) or non-threatening (non-aggressive) behavioural display (see **Figure 1**). For canines, aggressive displays specifically depicted offensive aggression to provide comparable threat-relevant postures across wolves and dogs. Crucially, to ensure that the displays predominantly engage implicit stimulus-driven responses, participants were only instructed to passively view the stimuli, without knowing they would later evaluate them. Only after completion of the passive viewing paradigm, participants were asked to rate each image for perceived threat, allowing us to relate neural responses to explicit threat evaluations. Targeted analyses of *a priori* selected subcortical regions assessed automatic threat responses and were complemented by exploratory whole-brain analyses of neocortical areas.

**Figure 1.**
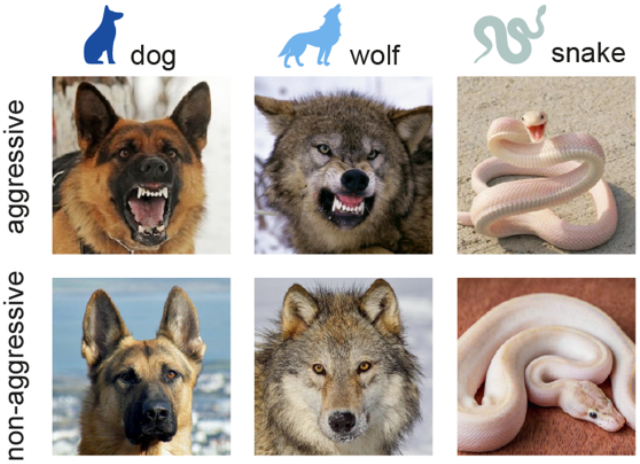
Experimental design. Participants saw images of wolves, dogs and snakes with an aggressive or non-aggressive behavioural display. Canine stimuli exclusively depicted offensive aggression; to match postures and perceived threat levels across canines, images were cropped to focus on the animals’ faces, and very small dog breeds were excluded. Participants first viewed the images while undergoing functional MRI, targeting their implicit neural responses, but after scanning explicitly assessed the perceived threat-levels of the depicted animals.

### Snakes evoke stronger subcortical threat responses than wolves or dogs

We first localised a functionally defined threat-responsive network using the aggressive > non-aggressive contrast pooled across all animals (see **Supplementary Figure S1**). This functional map was then intersected with a priori defined anatomical regions-of-interest (ROIs) in subcortical brain areas implicated in automatic threat detection, ensuring both high sensitivity and anatomical specificity. Because these ROIs were defined using the aggressive > non-aggressive contrast, extracting that same contrast for species comparisons would be circular; we therefore quantified implicit threat responses by extracting activation to aggressive displays of each species within these regions (i.e, central and basolateral amygdala, anterior and ventral insula.

The analysis revealed significant effects within the basolateral amygdala and the dorsal and ventral anterior insula, depending on the species viewed (**Figure 2A, Supplementary Table S1**). In all three ROIs, aggressive snakes elicited significantly stronger activation than aggressive wolves. Compared to dogs, snakes elicited greater activation only in the anterior insula, with no significant differences in the basolateral amygdala or ventral insula. By contrast, threat responses towards aggressive wolves vs. dogs significantly differed only in the basolateral amygdala, where dogs elicited stronger rather than weaker activation than wolves. We did not observe potential within-trial adaptation effects (e.g., faster vs. delayed neural responses to species) when conducting an exploratory time course analysis (see **Supplementary Figure S2**).

**Figure 2.**
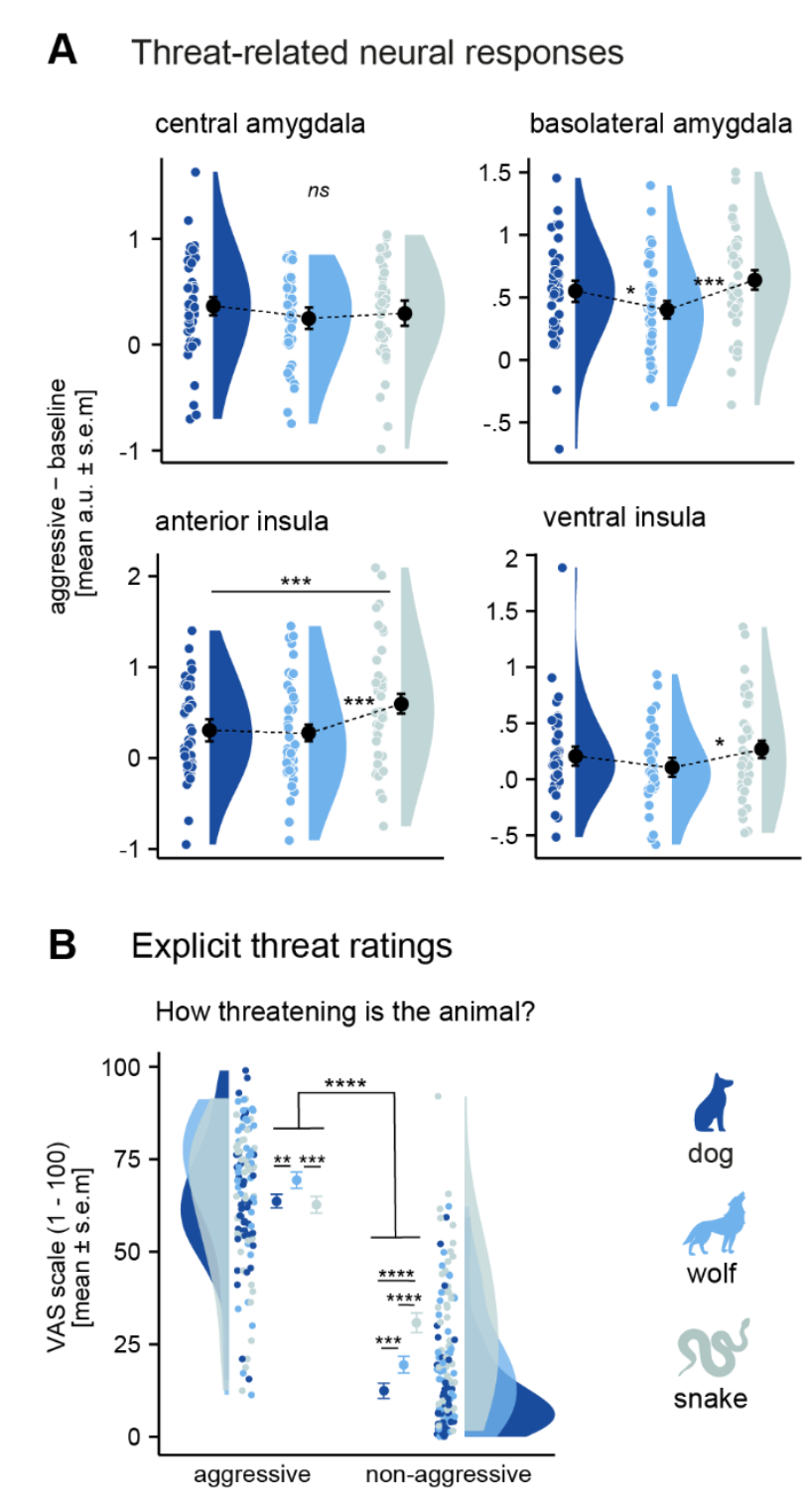
Divergent patterns of threat-related neural responses and explicit threat evaluations. (**A**) The functionally constrained region-of-interest analysis revealed greater threat responses for aggressive snakes than wolves in the basolateral amygdala and anterior and ventral insula. Compared to dogs, snakes elicited stronger activation only in the basolateral amygdala. Aggressive dogs elicited stronger activation than wolves in the basolateral amygdala, with no differences in the other regions (**Supplementary Table S4**) (**B**) Self-reported threat ratings (VAS) were higher for aggressive than non-aggressive displays, with aggressive wolves perceived as the greatest threat overall. Non-aggressive dogs were rated as the lowest threat, followed by non-aggressive wolves, then snakes. Aggressive dogs and snakes were rated as similarly threatening (see **Supplementary Table S3**). Raincloud plots (21) show group means (standard error of the mean, s.e.m.), individual means (dots), and distributions (half violins). \**p* <.05, \*\**p* <.01, \*\*\**p* <.001, \*\*\*\**p* <.0001; ns, not significant; a.u., arbitrary units.

### Explicit evaluations diverge from neural responses

Analysis of the explicit threat evaluations, measured via self-report after the MRI task, revealed a different pattern from the neural threat-related responses (**Figure 2B, Supplementary Table S2**). Threat evaluations differed for the species displayed, as indicated by a significant species × behavioural display interaction. Aggressive wolves were rated as the most threatening overall, exceeding all other species × display combinations and thus showing a pattern that diverged from the observed neural threat-related responses (cf. **Figure 2A**). Dogs were consistently rated as less threatening compared to their wolf counterparts, regardless of behavioural display. Compared with snakes, dogs were rated similarly in aggressive displays but less threatening in non-aggressive displays, where snakes received the highest ratings. Across all three species, perceived threat was higher for aggressive than for non-aggressive displays.

To examine the relationship between explicit threat evaluations and neural responses more directly, we conducted linear regression analyses testing the association between activation in response to aggressive displays of each species and the corresponding threat ratings for each ROI. None of them reached significance (see **Supplementary Table S3**).

### Whole-brain threat-related responses reveal widespread cortical activation and snake-specific subcortical engagement

To complement the ROI analyses and identify potential effects outside the predefined threat-related regions, we conducted exploratory whole-brain analyses. At the whole brain level, aggressive compared with non-aggressive behavioural displays of each species elicited greater activation in the primary visual cortex, as well as in higher-order occipito-temporal association areas, such as the lateral occipital complex (LOC) and the inferior and lateral temporal cortex (**Figure 3A, Supplementary Table S4**). For the same contrast, aggressive dogs and snakes additionally recruited parietal, premotor, and somatosensory regions, whereas only aggressive snakes engaged subcortical structures, including the right amygdala and bilateral insular cortex, consistent with the ROI results, which showed that snakes elicited the strongest subcortical threat-related responses (cf. **Figure 2B**). The reverse contrast (non-aggressive > aggressive) revealed greater activation in the hippocampus for dogs and in early visual areas for snakes, but no significant activation for wolves (**Figure 3B**). Together with the widespread activation of the aggressive > non-aggressive contrast, this pattern indicates that aggressive displays generally elicited stronger and more extensive whole-brain responses than non-aggressive displays, and this effect was observed uniformly across all species.

**Figure 3.**
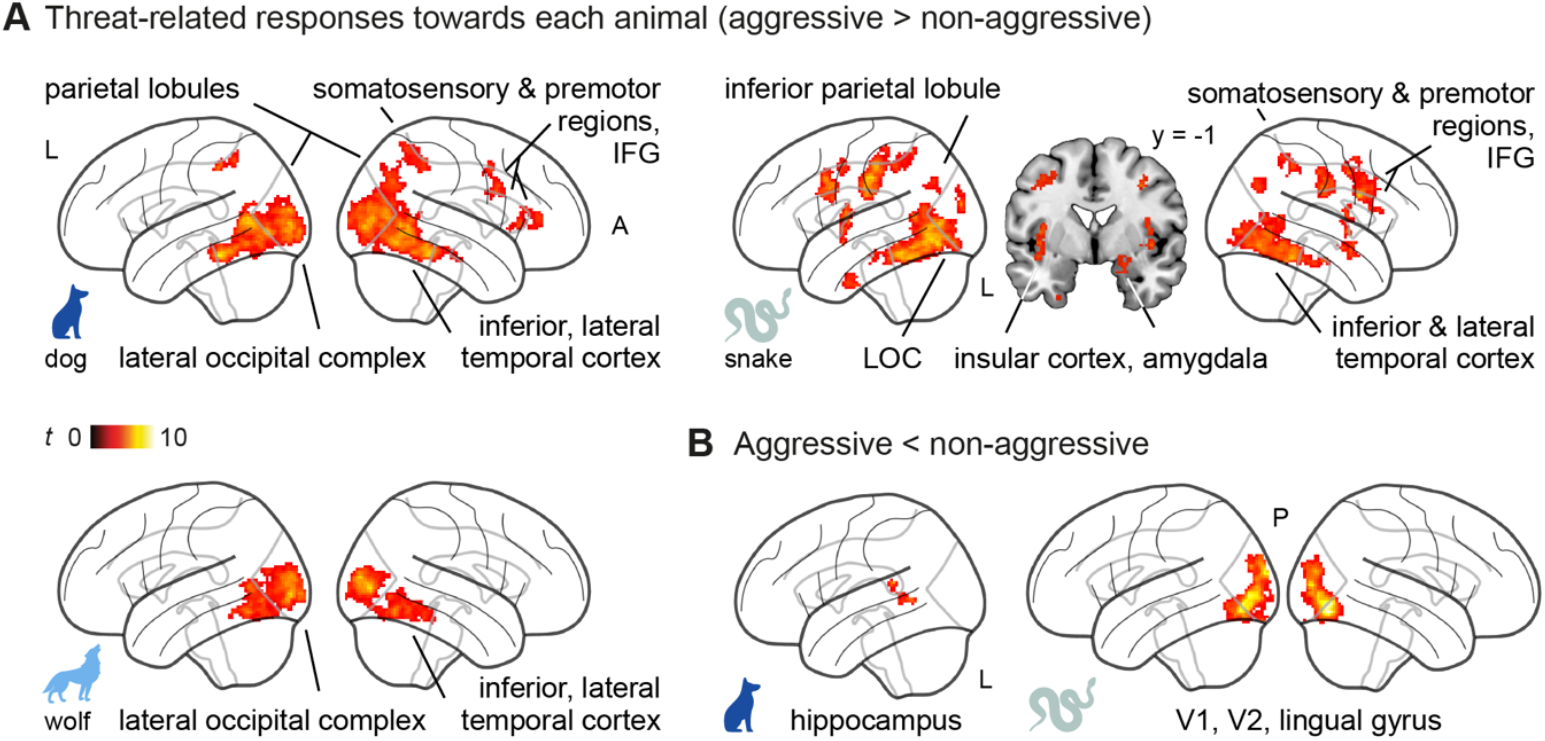
Whole-brain threat-related responses reveal widespread cortical activation and snake-specific subcortical engagement. (**A**) Compared to non-aggressive behavioural displays, aggressive displays led to greater activation in higher-order occipito-temporal association cortex for all animals. Viewing aggressive dogs or snakes additionally recruited partial and premotor regions, but only snakes (top right) show activation in subcortical limbic regions, including the amygdala and insular cortex. (**B**) For the reverse contrast (non-aggressive vs. aggressive) we found greater activation in the hippocampus and primary visual cortex when viewing non-aggressive dogs and snakes but no significant effects for non-aggressive wolves (see **Supplementary Table S4**). Results are *p* <.05 FWE-corrected at cluster-level using a cluster-defining threshold of *p* <.001. LOC, lateral occipital complex; V1/2, primary/secondary visual cortex; IFG, inferior frontal gyrus; A, anterior; P, posterior; R, L, left; *t*, t-values.

Whole-brain linear regression analyses then tested whether threat-related activation during aggressive displays covaried with the corresponding explicit threat ratings, but revealed no significant covariation between brain activation and perceived threat levels for any of the species. Likewise, no significant relationships were found for the difference between activation for aggressive wolves compared with aggressive dogs and the corresponding threat evaluations.

### Viewing dogs elicits higher-order visual cortex activation, while wolves elicit early visual cortex responses

After establishing the overall whole-brain threat-related responses for all animals, we focused on directly comparing neural responses to wolves and dogs across behavioural displays. This analysis revealed a significant main effect of behavioural display (aggressive > non-aggressive), with cortical activation patterns largely resembling the threat responses observed for each canine separately (**Figure 4A**, cf. **Figure 3A**). These included the lateral occipital and inferior temporal cortex, as well as parietal and somatosensory regions. We also found a main effect of species: irrespective of whether aggressive or not, viewing wolves recruited the primary visual cortex more strongly than viewing dogs, whereas dogs elicited greater activation in higher-order visual areas, including the lateral occipital complex and inferior temporal cortex (**Figure 4B**). Finally, a significant canine × behavioural display interaction revealed stronger activation in the inferior frontal and precentral gyrus for aggressive > non-aggressive dogs compared to the same contrast for wolves (**Figure 4C**).

**Figure 4.**
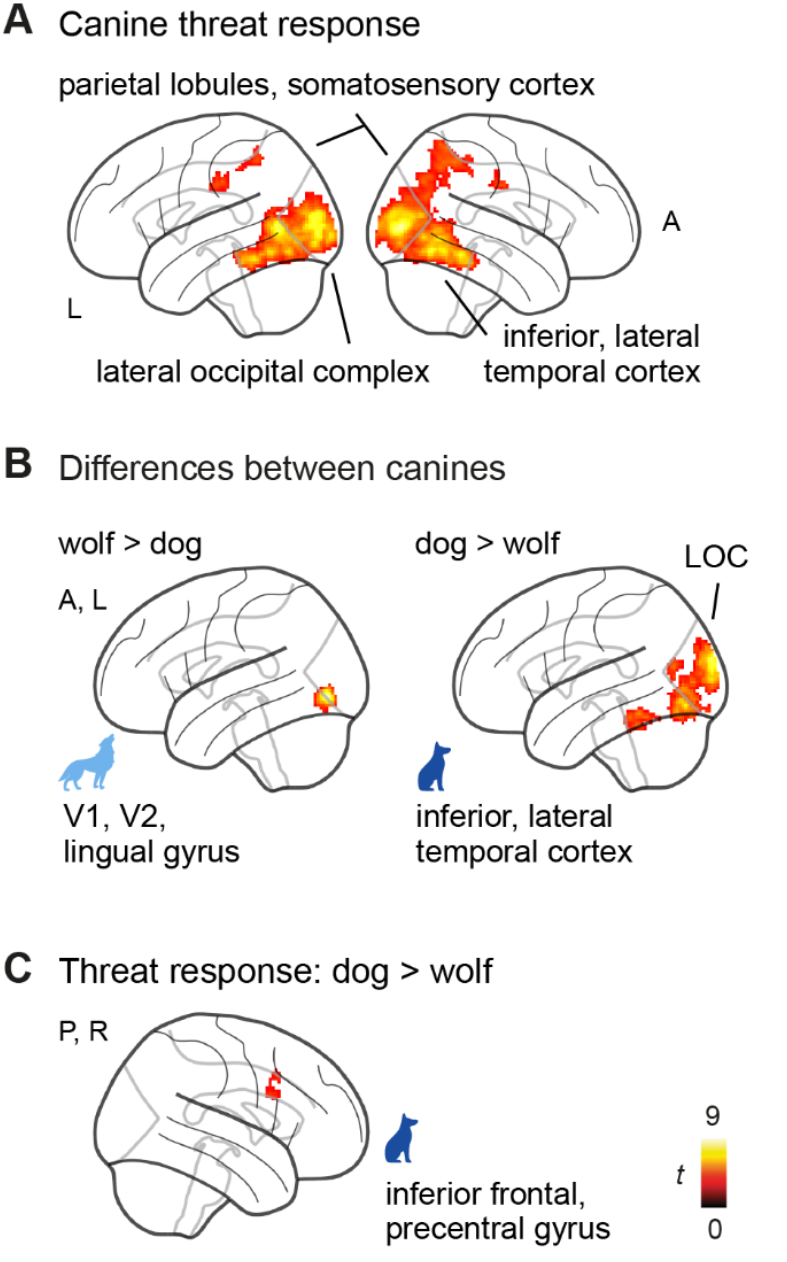
Viewing dogs elicits higher-order visual cortex activation, while wolves elicit early visual cortex responses. (**A**) Aggressive compared to non-aggressive behavioural displays led to activation in higher order parietal and occipito-temporal regions, as well as in the somatosensory cortex. (**B**) Viewing wolves compared to dogs regardless of affective display elicited significantly stronger activation in primary and secondary visual areas (V1, V2), the reversed contrast revealed significant activation in the lateral and inferior occipito-temporal cortex. (**C**) Aggressive compared with non-aggressive dogs elicited stronger activation in the inferior frontal and precentral gyri than did the corresponding contrast for wolves; see **Supplementary Table S5** for detailed results. LOC, lateral occipital complex; A, anterior; L, left; R, right; *t, t*-values.

## Discussion

The present study investigated the neural and behavioural bases of human attitudes toward wolves, their domestic counterparts, and snakes, using them as an established reference for evolutionarily prepared threat responses (15, 18, 19, 22). By combining functional neuroimaging with explicit threat ratings, we tested whether aversive perceptions of wolves reflect implicit-automatic, evolutionarily ingrained fear responses or rather explicit evaluative processes shaped by culture and experience. Our findings indicate that wolves, unlike snakes, do not elicit stronger subcortical threat responses than dogs, yet they were rated as the greatest threat. This dissociation suggests that opposition to wolves is unlikely to reflect evolutionary preparedness, but instead arises from evaluative processes shaping our perception of the ‘bad wolf’ and the ‘good dog’.

One of our main predictions was that if aversive attitudes toward wolves reflect an implicit, evolutionarily grounded threat bias, wolves would recruit subcortical brain areas canonically implicated in rapid and largely automatic threat detection, similar to those engaged by snakes. Contrary to our prediction, wolves did not elicit greater activation than dogs in these areas, neither in targeted region-of-interest (ROI) analyses nor in complementary whole-brain analyses. Across both approaches, and consistent with prior work, snakes showed the strongest engagement of subcortical threat-related brain areas. In the ROI analyses, activation differences for snakes were most pronounced in the basolateral amygdala and the anterior and ventral insula. The basolateral amygdala, closely interconnected with cortical regions, has been implicated in stimulus-specific and contextual evaluation of threats (23), while the anterior and ventral insula are involved in interoception, emotional valence assignment, and the regulation of anxious emotions (24–26). At the whole-brain level, the central amygdala, which is typically associated with immediate, visceral threat responses (15, 18, 19, 22, 23), also showed stronger activation for aggressive compared with non-aggressive snake displays, although this effect did not translate into greater activation for snakes relative to wolves or dogs in the ROI comparisons. Overall, these findings confirm previous observations of snakes as a highly salient threat, but they also emphasise that the human threat detection system is guided more by the immediate behavioural cues an animal displays than by its ancestral category.

In contrast to snakes, viewing wolves and dogs predominantly recruited cortical areas involved in higher-level visual processing, such as occipito-temporal regions for agent perception (27, 28), as well as parietal and premotor regions associated with action observation and execution (29–31). Dogs, compared to wolves, elicited greater activation in the lateral occipital complex and inferior temporal cortex regions associated with the perception of faces and bodies or social agents more generally (27, 28, 32). Wolves, in turn, elicited stronger responses in early visual areas, likely reflecting heightened visual attention (33). Together, these findings point to a qualitative difference in threat processing: Snakes triggered the strongest and most consistent responses in subcortical areas linked to the automatic threat detection and defensive circuitry, as described by dual systems accounts of fear and anxiety (20), whereas wolves and dogs elicited stronger engagement of cortical regions involved in reasoning, communication, and the evaluation of potential behavioural responses. Stronger activation for wolves in early visual areas may further reflect heightened vigilance toward uncertain or potentially threatening agents.

While neuroimaging revealed largely comparable threat processing for wolves and dogs, participants nonetheless rated wolves as more threatening, with aggressive wolves receiving the highest scores overall. Consistent with this dissociation, these explicit evaluations did not show any significant association with activation in the amygdala or insula, nor with whole-brain threat responses. Taken together with the finding that snakes elicited the strongest subcortical threat responses, this pattern suggests that negative views of wolves, and their frequent portrayal as dangerous, may arise less from automatic, evolutionarily ingrained fear mechanisms, but rather from socio-cultural influences shaping explicit evaluations.

### Limitations and future directions

Our study offers first insights into how implicit and explicit mechanisms shape human attitudes towards wolves. Several aspects should be considered when extending this work. First, the sample was recruited to be homogeneous in age, education, and cultural background to maximize experimental control. More diverse samples are needed to generalize our findings and to disentangle the contributions of cultural narratives, personal experience, and individual differences such as empathy or prior animal interactions. While fMRI can provide insights into where and how threatening stimuli are processed in the brain, it cannot fully capture implicit responses. Further studies, including complementary measures such as heart rate variability or skin conductance, as well as cognitive-behavioural measures of implicit processes, are thus needed to further disentangle early visceral responses from later evaluative processes. One potential avenue is, for example, the use of fear learning paradigms to test whether humans are predisposed to associate wolves with negative information compared to dogs, and how this is shaped by demographic or cultural factors (see e.g., 34 for a similar approach to study racial bias).

### Conclusion

Our results challenge the notion that humans are evolutionarily predisposed to fear wolves. While snakes elicited strong neural threat responses largely consistent with preparedness accounts, wolves did not, despite being rated as the greatest explicit threat. This dissociation indicates that aversive perceptions of wolves are unlikely to reflect automatic, evolutionarily ingrained threat detection mechanisms, but rather explicit evaluative processes shaped by knowledge, culture, and experience. This carries theoretical and practical implications. Theoretically, it underscores the flexibility of human fear systems and the importance of integrating cultural context into models of attitude formation. Practically, it suggests that negative attitudes toward wolves may be amenable to change: if rooted in explicit evaluation rather than in automatic neural biases, they should be more amenable to being shifted through education, communication, and carefully designed coexistence strategies. Recognizing that “bad wolves” are made rather than born in human minds may help enhance efforts toward human-wolf coexistence and, by extension, biodiversity conservation.

## Materials and Methods

### Participants

Forty-two participants (24 females; mean age: 23.86, *SD* = 2.43 years) participated in the functional MRI study. In the absence of prior studies and effect sizes, we determined the sample size based on previous studies in our lab with similar animal stimuli and task designs (27, 29). Participants were right-handed with normal or corrected-to-normal vision. They had no history of neurological or psychiatric diseases, did not report strong fear or phobia of dogs, wolves or snakes, and gave informed written consent. Participants were recruited through the Vienna CogSciHub: Study Participant Platform (SPP) using the hroot software (35). Data collection was approved by the ethics committee of the University of Vienna (reference number: 00565) and performed in line with the latest revision of the Declaration of Helsinki (2013).

### Experimental design

#### MRI task – implicit threat-related responses

In two 5-minute task runs, participants saw colour images of wolves, dogs, and snakes with aggressive or non-aggressive displays (see below for stimuli set details) on an MR-compatible 32-inch screen. We implemented the task as a passive-viewing paradigm. Crucially, participants were instructed to attend to the images presented on the screen, but did not know that they would have to assess the images’ threat level after completing the scanner task. We employed a block design (block duration: ∼10 s) with five images per block (2 s each). For each participant, image composition was randomized and each image was shown once per task run; block order was pseudo-randomized with no consecutive blocks of the same condition. The blocks were interspersed with a visual baseline (i.e., a white cross on a grey background) with a jittered intertrial interval of 3-7s, and the task was implemented using Psychopy (36).

#### Explicit threat assessment

After the MRI task, we asked participants to rate the perceived threat level of the images using a visual analogue scale (VAS) ranging from 0 (not threatening) to 100 (extremely threatening). Only one-third of the stimuli was rated as participants had reported fatigue when using the full stimuli set during piloting.

#### Stimulus material

The stimuli set consisted of 90 images, with 30 unique images for each species (wolf, dog, snake) representing various breeds or sub-species (see **Figure 1A** for examples). Half of the images depicted an aggressive display, while the other half depicted a non-aggressive display, matched by breed or sub-species respectively. We only selected dogs and wolves displaying offensive aggression displays and cropped the images to primarily display the canines’ faces to minimize potential effects of differences in size and posture. For the dog images, we included a variety of breeds but excluded very small ones (e.g., chihuahuas) to ensure comparable potential threat levels across canines. All images were standardized to 500 × 500 pixels to ensure consistent visual presentation.

### MRI data acquisition and preprocessing

We acquired the MRI data with a 3T Siemens Skyra MR system (Siemens Medical, Erlangen, Germany) using a 32-channel head coil. Functional scans were obtained with a 4-fold multiband accelerated EPI sequence: voxel size = 2 mm isotropic, repetition time (TR) / echo time (TE) = 1200/34 ms, FoV = 192 × 192 × 124.8 mm^3^, flip angle = 66°, 20% gap and 52 axial slices coplanar to the connecting line between anterior and posterior commissure (interleaved acquisition, ascending order). In the same orientation as functional scans, we obtained additional field map scans to correct for magnetic field inhomogeneities using a double echo gradient echo sequence with a voxel size of 1.72 × 1.72 × 3.85 mm^3^, TR/TE1/TE2 = 400/4.92/7.38 ms, FoV = 220 × 220 × 138 mm^3^, flip angle = 60° containing 36 axial slices. Structural images were acquired using a MPRAGE sequence with the following parameters: voxel size =.8 mm isotropic, TR/TE = 2300/2.43 ms and FoV = 256 × 256 × 166 mm^3^. We preprocessed the neuroimaging data using standard SPM12 (https://www.fil.ion.ucl.ac.uk/spm/software/spm12/) pipelines (see **Supplementary material**).

### Data analysis overview

We investigated implicit and explicit processes underlying attitudes toward wolves and dogs, with snakes included as a reference condition for evolutionarily prepared threat responses. Analyses were conducted at two complementary levels. A targeted functional region-of-interest (fROI) approach tested *a priori* hypotheses concerning subcortical brain regions implicated in automatic threat processing, while exploratory whole-brain analyses investigated additional threat-related activation patterns beyond the ROI-defined brain areas. Explicit threat evaluations (self-reports) for all animals and behavioural displays were analysed and subsequently related to corresponding neural responses through ROI-wise and whole-brain regression analyses.

The fROI analysis assessed differences in mean activation levels and, in supplementary analyses, the temporal dynamics of activation levels during aggressive displays. Exploratory whole-brain analyses investigated activation for aggressive versus non-aggressive displays of each species to characterize general threat-related activation patterns and directly compared neural responses to wolves and dogs as a function of behavioural display to isolate species-specific effects, consistent with the main focus of the study.

### MRI analysis

#### First-level analysis

For the individual first-level GLM design matrices, we defined task regressors for the six conditions of interest: the aggressive and non-aggressive behavioural display of each species (i.e., dog/wolf/snake × aggressive/non-aggressive display). All blocks were estimated as a boxcar function time-locked to the onset of each block with the respective durations and convolved with the canonical HRF. The six motion regressors retrieved from image realignment and the individual framewise displacement regressors were added nuisance regressors, and we applied a temporal high-pass filter with a cut-off at 128 s. We then computed contrasts for all six task regressors, a threat response contrast for each animal (i.e., aggressive > non-aggressive), and a general threat response pooled across all animals (i.e., aggressive > non-aggressive).

#### Functional region-of-interest (ROI)

Based on prior research on threat perception, we expected the central (CeA) and basolateral (BLA) amygdala (20, 23), the anterior (aINS) and ventral (vINS) insular cortex (24–26) and bed nucleus stria terminalis (BNST; 37) to play key roles in implicit threat encoding. Anatomical masks of the CeA, BLA and the BNST were obtained from existing parcellations (37, 38). The aINS mask was created by combining areas Id7 and Id6, and the vINS mask by merging Id8 and Id10 of the Julich-Brain atlas (39). These anatomical masks were then intersected with the pooled whole-brain threat response (aggressive > non-aggressive across animals) to extract and analyse (i) mean activation levels, (ii) the temporal response dynamics, and (iii) brain-behaviour associations. Using an orthogonal contrast for intersection maximized sensitivity while maintaining anatomical specificity and ensured that the ROIs remained statistically independent of the contrasts of interest, thereby avoiding circularity. The BNST had to be removed from the analysis due to a lack of overlap between the anatomical mask and the pooled whole-brain threat response. *P*-values for planned post hoc and group comparisons investigating the same research questions were FDR-controlled.

#### Mean activation levels

We extracted parameter estimates (i.e., activation levels) using the REX toolbox (40). To assess threat-related responses, we conducted linear mixed-effects models (LMMs) separately for each of the four ROIs, using the R packages *lme4* (41) and *afex* (42). In each model, activation levels for aggressive displays served as the dependent variable, *species* (dog, wolf, snake) as the predictor, and subject as a random intercept.

#### Mean time course of activation

Even in the absence of differences in activation levels averaged across the entire stimulus presentation, threat responses might still differ over time, particularly given the block design of our study. To test this possibility, we set up supplementary analyses (see **Supplementary material** for details).

#### Behavioural data analysis and brain-behaviour relationship

For the analysis of explicit behavioural threat assessments, we employed a LMM with threat ratings as the dependent variable. We defined species (dog, wolf, snake) and *behavioural display* (aggressive, non-aggressive) as predictors and added per-subject random intercepts.

To examine how individual differences in explicit threat evaluations predicted neural activation, we conducted ROI-based regression analyses. For each ROI, mean activation levels for aggressive displays of each species (dogs, wolves, snakes) served as the dependent variable, with the corresponding threat ratings as predictor. Additionally, to assess whether inter-individual differences in *relative* threat evaluation corresponded to *relative* neural activation, we performed ROI-wise regression analyses using difference scores (aggressive wolves – aggressive dogs) for activation (dependent variable) and threat ratings (predictor).

#### Whole-brain group comparisons

For the complementary whole-brain analysis, we conducted one sample *t*-tests for each threat response and the reversed contrast to explore whole-brain activation towards each animal. We then focused on the comparison between our main conditions of interest and calculated a 2-by-2 within-subjects full factorial model using the flexible factorial framework in SPM12 with factors *canine* (wolf, dog) and *behavioural display* (aggressive, non-aggressive), and a canine × behavioural display interaction. We also conducted a one-sample *t*-test for the pooled threat response thresholded at *p* =.005 to serve as a functional mask for the above-described functional ROI analyses. A liberal threshold is standard for functional localizers because the resulting mask is used solely to define the search space, rather than for statistical inference.

#### Whole-brain linear regression analysis

Mirroring the ROI-level analysis, we then employed multiple regression analyses for the contrasts dog/wolf/snake aggressive > implicit baseline and aggressive wolf > aggressive dog, in which we added the corresponding threat ratings as covariates.

For all group comparisons, we determined significance by applying cluster-level inference with a cluster-defining threshold of *p* <.001 and a cluster probability of *p* <.05 family-wise error (FWE) corrected for multiple comparisons and derived the cluster extent (i.e., the minimum spatial extent to be labelled significant) using the SPM extension “CorrClusTh.m”. Anatomical labelling of activation peaks and clusters of all reported results was performed using the python software AtlasReader (43), and labels refer to the Julich-Brain atlas (44) and the Harvard-Oxford brain atlas (45).

## Data and code availability

We analysed the data using Matlab 2020b (MathWorks), SPM12 (https://www.fil.ion.ucl.ac.uk/spm/software/spm12/), and R 4.4.0. We employed all linear mixed models using the R packages *lme4* (41) and *afex* (42), and created the corresponding figures with *ggplot2* (46) and *RainCloudPlots* (21). Whole-brain neuroimaging figures were created with the python project nilearn (http://nilearn.github.io) and MRIcron (https://www.nitrc.org/projects/mricron). The task was implemented using PsychoPy (36). Individual rating and ROI data, as well as whole-brain statistical maps, are openly available on the Open Science Framework (OSF; osf.io/v6ajq).

## Author contributions (CRediT)

**M.B**.: Conceptualization, Investigation, Data curation, Methodology, Software, Formal analysis, Writing – original draft, Writing – review & editing, Visualization, Project administration. **F.R**.: Conceptualization, Writing – review & editing, Funding acquisition. **S.M-P**.: Conceptualization, Writing – review & editing. **R.S**.: Formal analysis, Methodology, Writing – review & editing. **C.L**.: Conceptualization, Resources, Writing – original draft, Writing – review & editing, Supervision, Funding acquisition.

## Acknowledgements

We want to thank Anna Thallinger, Olaf Borghi and Sara Binder for their support in collecting the data for this project.

