## Supplementary material for "Good dog, bad wolf: Debunking evolutionary preparedness to fear wolves"

### 1 Supplementary notes and figures

#### Extended Materials and Methods

**MRI preprocessing pipeline.** We preprocessed and analyzed the neuroimaging data using SPM12 (https://www.fil.ion.ucl.ac.uk/spm/software/spm12/) and Matlab 2020b (MathWorks Inc., MA, USA). After performing slice timing correction using the middle slice as a reference (1), we realigned and unwarped the functional scans using the individual field maps. Next, the individual structural scans were co-registered to the mean functional image and segmented. Finally, we registered the functional scans to MNI space via the co-registered structural scan and applied spatial smoothing with a 3-dimensional Gaussian Kernel (FWHM, 4 mm, i.e., twice the raw voxel resolution). Based on the six realignment parameters, we calculated individual scan-to-scan motion (i.e., framewise displacement, FD), and for each scan exceeding the FD threshold of .3 mm, we added a motion regressor to the first-level general linear models (2, 3). On average, 7% of the scans exceeded the threshold in each run (run 1: mean FD = .17 mm, 90$^{th}$ percentile = .21 mm; run 2: mean FD = .17 mm, 90$^{th}$ percentile = .22 mm).

**Individual finite impulse response models.** For supplementary analyses, we also set up individual finite impulse response (FIR) models to estimate the time course of activation for all six conditions of interest. Per condition, we estimated FIRs covering a time window of 12 s starting 5 s after block onset (block duration = 10 s). The 12 s were then divided into 12 time bins (i.e., one time bin per TR) and modelled with separate regressors using an impulse response function. We then extracted the average BOLD signal time course in areas hypothesized to encode threat responses (see **section ROI analyses**).

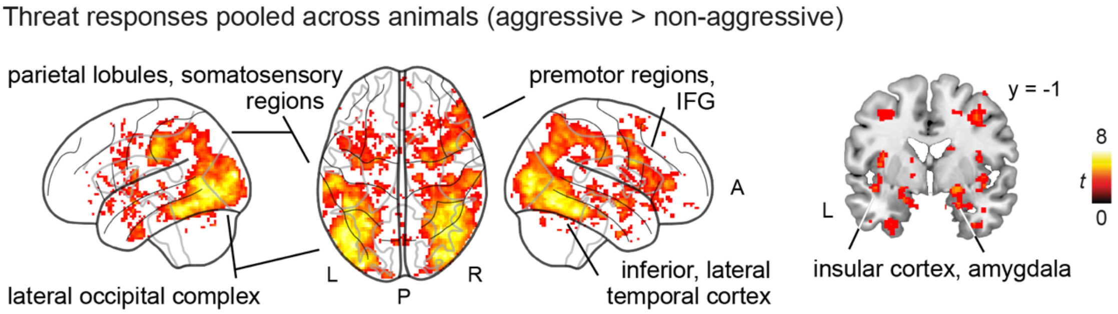


**Supplementary Figure S1. Pooled threat response across all animals to serve as functional mask for region-of-interest analysis.** Aggressive displays pooled across all animals led to distributed whole-brain activation including temporo-parietal and frontal cortex as well as subcortical brain regions. Results are liberally thresholded at $p$ = .005 uncorrected. The intersection between the functional results and anatomical masks served as regions-of-interest (ROI) to compare threat response between animals.

Next, we extracted the estimated time courses for regions of interest and visualized them over time. We did not find evidence for a systematic difference in the activation profile in the amygdala (central and basolateral) or the insula (anterior, ventral), other than the higher activation for the snake stimuli. As a control, we also plotted the time course of the visual cortex, where we observed a strong BOLD response to all stimuli (see **Supplementary Figure S2**).

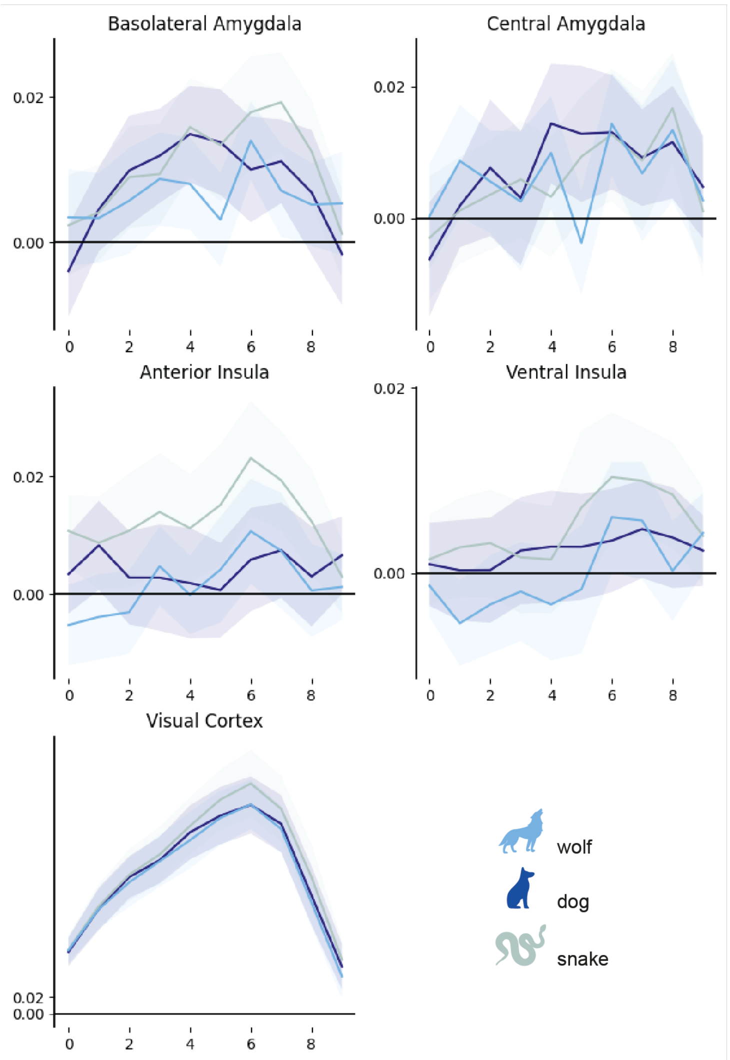


**Supplementary Figure S2. Time course of the ROIs as estimated by the FIR model.** X-axis: Time bin (seconds); Y-axis: beta (arbitrary units); Shaded area: 95% confidence interval. Dog, dark blue; Wolf, light blue; Snake, teal.

### 2 Supplementary tables

**Table S1.** Functional regions-of-interest analysis LMM results[a]

| Region-of-interest | F ($df_{Num}$, $df_{Den}$) | $p$ | $p_{FDR}$ | $\eta^2_g$ |
|---|---|---|---|---|
| Central amygdala | 1.25 (2,81) | .2918 | .2918 | .03 |
| Basolateral amygdala | 9.76 (2,81) | **.0002** | **.0004** | .19 |
| Anterior insula | (2,81) | **.0001** | **.0004** | .21 |
| Ventral insula | (2,81) | **.0221** | **.0295** | .09 |

*Note.* [a]Within-subjects design, dependent variable: activation levels for aggressive behavioural displays, predictor: *animal* (levels: dog, wolf, snake) with a random intercept for each subject. The model was estimated using restricted maximum likelihood. Functional regions-of-interest (ROIs) were created by intersecting *a priori* defined subcortical regions-of-interest (ROIs) with functional thread localizer contrast (aggressive > non-aggressive pooled across animals; see **Supplementary Figure S1C**). *P*-values for group comparisons are false discovery rate (FDR) corrected as they investigate the same research question in each ROI; uncorrected *p*-values are also reported, and *p*-values < .05 are in bold. Results of post hoc comparisons are displayed in **Figure 2A**. $df_{Num}$, degrees of freedom numerator; $df_{Den}$ degrees of freedom denominator; $\eta^2_g$, generalized eta-squared.

**Table S2.** Explicit threat ratings LMM results[a]

| Self-report threat ratings | F ($df_{Num}$, $df_{Den}$) | $p$ | $\eta^2_g$ |
|---|---|---|---|
| Animal | 22.52 (2, 1211) | **< .0001** | .04 |
| Behavioural display | 1626.03 (1, 1211) | **< .0001** | .57 |
| Animal × behavioural display | 32.18 (2, 1211) | **< .0001** | .05 |

*Note.* [a]2 × 3 within-subjects design, dependent variable: explicit threat ratings, predictors: *animal* (levels: dog, wolf, snake) and *behavioural display* (aggressive, non-aggressive) with a random intercept for each subject. The model was estimated using restricted maximum likelihood; *p*-values < .05 are in bold. Results are displayed in **Figure 2B**; $df_{Num}$, degrees of freedom numerator; $df_{Den}$ degrees of freedom denominator; $\eta^2_g$, generalized eta-squared.

**Table S3.** ROI-wise linear regression analyses for brain-behaviour associations[a]

| Region-of-interest, contrast | β | *t* | *p* | $p_{FDR}$ |
|---|---|---|---|---|
| Central amygdala | | | | |
| Aggressive dogs | -.004 | -.89 | .3804 | .9739 |
| Aggressive wolves | -.003 | -.76 | .4494 | .9739 |
| Aggressive snakes | .003 | .91 | .3664 | .9739 |
| Aggressive wolves - aggressive dogs | -.0052 | -1.2 | .236 | .289 |
| Basolateral amygdala | | | | |
| Aggressive dogs | -.002 | -.6 | .5512 | .9739 |
| Aggressive wolves | .001 | -.29 | .7758 | .9739 |
| Aggressive snakes | .002 | .52 | .6097 | .9739 |
| Aggressive wolves - aggressive dogs | -.0043 | -1.08 | .289 | .289 |
| Anterior insula | | | | |
| Aggressive dogs | 0 | -.03 | .9739 | .9739 |
| Aggressive wolves | .002 | .42 | .6803 | .9739 |
| Aggressive snakes | .004 | .77 | .4439 | .9739 |
| Aggressive wolves - aggressive dogs | .0132 | 2.56 | **.0145** | .0579 |
| Ventral insula | | | | |
| Aggressive dogs | -.002 | -.67 | .5037 | .9739 |
| Aggressive wolves | 0 | .1 | .9242 | .9739 |
| Aggressive snakes | 0 | -.05 | .9566 | .9739 |
| Aggressive wolves - aggressive dogs | .005 | 1.14 | .262 | .289 |

*Note.* [a]Dependent variable: activation levels, predictor: corresponding explicit threat ratings through self-reports. Analyses were conducted per region-of-interest and contrast. *P*-values for group comparisons are false discovery rate (FDR) corrected as they investigate the same research question in each ROI; uncorrected *p*-values are also reported, and *p*-values < .05 are in bold- *t*, *t*-values.

**Table S4.** Whole-brain threat responses towards each animal

| Contrast & brain region | coordinates | | | *z*-value | cluster size |
|---|---|---|---|---|---|
| | x | y | z | | |
| **Aggressive wolves > non-aggressive wolves** (*k* = 55) | | | | | |
| R lateral occipital cortex, inferior division | 32 | -84 | 4 | 5.65 | 1709 |
| L occipital pole | -32 | -94 | 0 | 5.55 | 2205 |
| **Aggressive dogs > non-aggressive dogs** (*k* = 54) | | | | | |
| L lateral occipital cortex, inferior division | -44 | -40 | -20 | 5.88 | 3127 |
| R lateral occipital cortex, inferior division | 46 | -40 | -18 | 5.47 | 4479 |
| L superior parietal lobule | -26 | -40 | 46 | 4.61 | 79 |
| R inferior frontal gyrus pars triangularis | 52 | 28 | 4 | 4.28 | 169 |
| R superior parietal lobule | 28 | -44 | 50 | 4.23 | 288 |
| R precentral gyrus | 50 | 12 | 32 | 4.17 | 150 |
| R middle frontal gyrus | 34 | 6 | 44 | 3.99 | 55 |
| **Aggressive snakes > non-aggressive snakes** (*k* = 51) | | | | | |
| L postcentral gyrus, inferior parietal lobule | -58 | -24 | 38 | 5.64 | 960 |
| L lateral occipital cortex, inferior division | -44 | -72 | -10 | 5.63 | 2559 |
| R precentral gyrus | 44 | 6 | 26 | 5.42 | 625 |
| R inferior temporal gyrus, temporooccipital part | 50 | -44 | -22 | 5.21 | 1969 |
| L precentral gyrus | -52 | 8 | 30 | 5 | 326 |
| L lateral occipital cortex, superior division | -40 | -90 | 18 | 4.99 | 66 |
| L insular cortex | -40 | -4 | 8 | 4.86 | 128 |
| R supramarginal gyrus, superior division | 64 | -20 | 32 | 4.65 | 328 |
| R insular cortex | 42 | -2 | -2 | 4.52 | 69 |
| R amygdala | 24 | -6 | -14 | 4.38 | 96 |
| L precentral gyrus | -42 | 0 | 44 | 4.22 | 86 |
| R superior parietal lobule | 30 | -44 | 48 | 4.21 | 96 |
| L temporal fusiform cortex, anterior division | -26 | -6 | -40 | 4.12 | 95 |
| R lateral occipital cortex, superior division | 28 | -70 | 32 | 4.11 | 60 |
| L lateral occipital cortex, superior division | -24 | -70 | 36 | 3.9 | 55 |
| **Aggressive dogs < non-aggressive dogs** (*k* = 54) | | | | | |
| L hippocampus | -20 | -40 | 16 | 5.04 | 74 |
| R hippocampus | 24 | -42 | 8 | 4.49 | 70 |
| **Aggressive snakes < non-aggressive snakes** (*k* = 51) | | | | | |
| R lingual gyrus | 10 | -78 | -12 | 7.17 | 2743 |

*Note.* Effects were tested for significance with a cluster defining threshold of $p$ <.001 and a cluster probability threshold of $p$ < .05 FWE. We report the first local maximum within each cluster for each $t$-test along with the critical cluster sizes ($k$). The contrast aggressive wolves < non-aggressive wolves did not result in any significant activation. The data is presented in **Figure 3** and anatomical labels refer to the Julich-Brain atlas (4) and the Harvard-Oxford brain atlas (5). L, left; R, right.

**Table S5.** Whole brain results from 2 × 2 full factorial model[a] to compare response towards canines

| Contrast & brain region | coordinates | | | *z*-value | cluster size |
|---|---|---|---|---|---|
| | x | y | z | | |
| **Aggressive > non-aggressive** | | | | | |
| R lateral occipital cortex, inferior division | 32 | -86 | 4 | 7.38 | 5593 |
| L lateral occipital cortex, inferior division | -50 | -64 | 6 | 6.8 | 4441 |
| L inferior parietal lobule | -64 | -26 | 36 | 4.33 | 129 |
| R postcentral gyrus | 56 | -14 | 34 | 4.16 | 74 |
| L superior parietal lobule | -24 | -50 | 50 | 4.09 | 62 |
| **Wolf > dog** | | | | | |
| L primary visual cortex (V1) | 0 | -82 | -6 | Inf | 374 |
| **Dog > wolf** | | | | | |
| L occipital pole | -12 | -98 | 18 | 6.79 | 5095 |
| L fusiform cortex | -40 | -48 | -26 | 4.91 | 223 |
| **Dog [aggressive > non-aggressive] > wolf [aggressive > non-aggressive]** | | | | | |
| R inferior frontal gyrus, pars opercularis | 36 | 18 | 40 | 3.88 | 131 |

*Note.* [a] Dependent variable: whole-brain activation levels, predictors: *animal* (levels: dog, wolf, snake) and *behavioural display* (aggressive, non-aggressive) with a random intercept for each subject. The model was estimated using restricted maximum likelihood. Effects were tested for significance with a cluster defining threshold of $p$ <.001 and a cluster probability threshold of $p$ < .05 FWE; the critical cluster sizes to determine significance was $k$ = 59 We report the first local maximum within each cluster for each $t$-test along with the critical cluster sizes ($k$). Post hoc comparisons for the contrasts non-aggressive > aggressive, wolf [aggressive > non-aggressive] > dog [aggressive > non-aggressive] did not survive multiple comparison corrections. The data is presented in **Figure 4** and anatomical labels refer to the Julich-Brain atlas (4) and the Harvard-Oxford brain atlas (5). L, left; R, right
